# Assessment of two immunoassays for detection of IgM antibodies to scrub typhus using a serum panel

**DOI:** 10.1101/667410

**Authors:** Divyaa Elangovan, Susmitha Perumalla, Winsley Rose, Valsan Philip Verghese, M.S. Gowri, John Antony Jude Prakash

## Abstract

**Introduction:** Scrub typhus is a vector borne zoonotic disease caused by *Orientia tsutsugamushi*, endemic to tsutsugamushi triangle. As the characteristic eschar is not always present, laboratory testing especially serological assay are the main stay of diagnosis.

**Materials and methods:** A total of 346 well-characterized sera from normals and patients with scrub typhus, malaria, dengue, enteric fever and gram negative septicaemia were tested for IgM antibodies by ST IgM ELISA and ST Ig M ICT

**Results:** The sensitivity and specificity of Scrub typhus IgM ICT and ELISA were 98.7, 96.3 and 97.4, 99.3 respectively. The IgM ICT and ELISA had a excellent concordance (99%) and a very high negative predictive value.

**Conclusion:** The findings from this study suggest that IgM ICT and IgM ELISA can be used interchangeably for serodiagnosis of scrub typhus in resource poor settings.

## Introduction

Scrub typhus is a mite-borne zoonotic disease caused by *Orientia tsutsugamushi*, endemic to South Asia, Southeast Asia, East Asia, the Pacific Islands, and Northern Australia; also called the “tsutsugamushi triangle”.^1,2^ Serological evidence from places in Africa, France, the Middle East and South America suggests that scrub typhus is no longer limited to the tsutsugamushi triangle.^3^ Scrub typhus (ST) is being increasingly reported from all over India with a prevalence of over 30-40% of all acute febrile illness (AFI) cases needing admission to hospitals.^4,5^ Differentiating scrub typhus from other endemic causes of AFI based on clinical features alone is a daunting task for the clinician. Though the presence of an eschar at the mite inoculation site is a valuable diagnostic clue, it is not always seen in all patients. Additionally, the lack of wide availability of confirmatory tests augments problems with diagnosis. The combination of an eschar and a positive result on a rapid diagnostic test (RDT) is shown to have positive and negative predictive values of 84.9 and 93.0% respectively, but the varying occurrence of eschar limits this approach.^6^ Data from our tertiary care centre in South India has documented eschar rates close to 60%.^7,8^ However, the absence of an eschar does not rule out scrub typhus and hence diagnosis relies heavily on laboratory tests, especially serological tests. ^7^ The tests available for the diagnosis of ST come with their own limitations. In our experience, the inexpensive Weil-Felix serological test showed good specificity,^9^ but lacked sensitivity.^10^ Immunofluorescence assay is the serological reference standard for diagnosis of ST but its cost, availability only in reference labs and subjectivity in interpretation of results limit its use. IgM ELISA test which is objective has good sensitivity and specificity and can be automated allowing testing of large number of samples.^11^ Diagnosis using molecular methods is not available in secondary level health care settings like district hospitals in India. ELISA is easy and relatively economical, but samples need to be batch tested, which may delay the diagnosis and affect the overall disease outcome. It is evident that many of these laboratory tests cannot be accessed in rural settings where most of the cases occur.

With due consideration to expense, rapidity and simplicity of interpretation, rapid diagnostic tests have emerged. Although DHR-ICMR guidelines 2015 for the diagnosis and management of rickettsial diseases in India discourage the use of rapid tests, they do emphasize the need for further evaluation of rapid diagnostic kits.^12^ The use of immunochromatographic technology in development of RDTs offer a point-of-care serological test at bedside, which ultimately helps the clinician to start early management with the anti-rickettsial therapy especially in those with atypical presentation, thereby reducing morbidity and mortality.^8^

In this study, we evaluated two tests, an ELISA and an immunochromatographic test (ICT) which is the rapid diagnostic test (RDT) for detection of IgM antibodies to *Orientia tsutsugamushi* strain. Both these assays use recombinant *Orientia tsutsugamushi* 56-kD type-specific antigen from strains Kato, Karp, Gilliam, and TA716.^13^

## Materials & Methods

Serum samples sent for routine microbiology testing in our lab from 2016-2018 were stored as a part of serum banking protocol (IRB Min NO 7768). A total of 346 samples were tested for IgM antibodies by ELISA (Scrub typhus Detect IgM ELISA, InBios International Inc., Seattle WA, USA) as per already established protocol.^14^ Immunochromotography test to detect IgM antibodies for scrub typhus (Scrub Typhus Detect IgM rapid test, InBios International Inc., Seattle WA, USA) was performed as per the manufacturer’s instruction. Clinical details of patients were collected from the records of the patients made available on the electronic database of the study center.

Details of the samples used to assess the performance characteristics of both the tests are given in table 1

**Table 1:** Details of sera used for evaluation of ST IgM ELISA and ST IgM ICT.

| Clinical spectrum | Criteria used for classification of tested samples | Number tested |
| --- | --- | --- |
| Normal | No h/o fever for $\geq 3$ months | 71 |
| Dengue | NS1 Ag positive<br>(DENGUE NS1 Ag MICROLISA<br>J.Mitra & Co. Pvt. Ltd,<br>New Delhi, India) | 50 |
|  | Dengue IgM & IgG positive<br>(J.Mitra & Co. Pvt. Ltd,<br>New Delhi, India) | 8 |
| Enteric fever | Blood culture positive<br>(BacT/Alert 3D, Biomerieux,<br>Marcy l'Etoile, France) | 47 |
| Gram negative septicemia |  | 54 |
| Vivax malaria | Peripheral blood thick & thin smear positive for malarial and filarial parasites | 19 |
| Falciparum malaria |  | 19 |
| Microfilaria |  | 1 |
| ST cases * | IgM ELISA positive ± eschar; Fever responded to doxycycline within 48 hours | 77 |

\*Individuals with acute febrile illness, duration of illness $\geq 5$ days, malaria smear negative, blood culture and dengue negative

## Statistical Methods

Categorical data was expressed as numbers and percentages. The agreement between ICT and ELISA was presented with standard error (SE) and concordance rate. Diagnostic accuracies (sensitivity, specificity, LR+, LR-, PPV and NPV) were presented with 95% CI. Receiver operator characteristics (ROC) curve was constructed for OD values to discriminate the diseased and the non-diseased. The result of ROC is shown with AUC (95% CI) and the optimal cut-off along with sensitivity and specificity are described. All the analysis was done using STATA IC/15.1 (Stata Corp LLC, College Station, Texas, USA)

## Results

Among the 77 patients with ST positive serum samples, 51 had a characteristic eschar. ST IgM ELISA and ST IgM ICT demonstrated 11 and 9 false-positive results respectively, the details of which are enumerated in table 2.

**Table 2:** Details of positive results for scrub typhus.

| Clinical spectrum | Numbers tested | Positive ST IgM ELISA | Positive ST IgM ICT |
| --- | --- | --- | --- |
| Normal | 71 | 0 | 0 |
| Dengue | 58 | 6 | 6 |
| Enteric fever | 47 | 0 | 0 |
| Gram negative septicaemia | 54 | 0 | 0 |
| Vivax malaria | 19 | 2 | 1 |
| Falciparum malaria | 19 | 3 | 2 |
| Microfilaria | 1 | 0 | 0 |
| ST cases * | 77 | 77 | 77 |

\*Individuals with acute febrile illness, duration of illness $\geq 5$ days, malaria smear
negative, blood culture and dengue negative

All six dengue sera which were also positive in both ELISA and ICT tests were positive for NS1 antigen of dengue. A breakdown of all the false positive results on ELISA, ICT and the final diagnosis made are represented in table 3.

**Table 3:** Breakdown of the false positive results on ELISA and ICT.

| **Case number** | **Other positive parameter** | **ST IgM ELISA OD** | **ST IgM ICT** | **Fever duration** | **Final diagnosis** |
| --- | --- | --- | --- | --- | --- |
| **1** | NS 1 Ag ELISA | 3.04 | Positive | 10 days | Scrub typhus |
| **2** | NS 1 Ag ELISA | 2.55 | Positive | 4 days | Scrub typhus |
| **3** | NS 1 Ag ELISA | 2.88 | Positive | 15 days | Scrub typhus |
| **4** | NS 1 Ag ELISA | 2.55 | Positive | 7 days | Scrub typhus |
| **5** | NS 1 Ag ELISA | 1.89 | Positive | 7 days | Dengue |
| **6** | NS 1 Ag ELISA | 1.02 | Positive | 2 days | Dengue |
| **7** | Plasmodium falciparum | 1.48 | Positive | 7 days | Malaria |
| **8** | Plasmodium falciparum | 1.09 | Positive | ----- | Malaria |
| **9** | Plasmodium vivax | 1.07 | Positive | ----- | Malaria |
| **10** | Plasmodium falciparum- | 2.49 | Negative | 10 days | Malaria |
| **11** | Plasmodium vivax | 1.21 | Negative | ----- | Malaria |

Among the six patients with false positive results on dengue sera, a final diagnosis of ST was made in four based on clinical findings and response to ST-specific therapy, while the other two were considered false positive results

When agreement between IgM ICT and IgM ELISA was studied, almost perfect agreement between their results with a kappa value 0.97 in accordance with the agreement criteria ^15^ was found and the same is represented in table 4.

**Table 4:** Agreement between IgM ICT and IgM ELISA for scrub typhus diagnosis.

| Test done |  | ST IgM ICT |  | Concordance | Kappa(SE) |
| --- | --- | --- | --- | --- | --- |
|  |  | Neg | Pos |  |  |
| ST IgM ELISA | Neg | 258 | 2 | 98.8% | 0.97 (0.054) |
|  | Pos | 2 | 84 |  |  |

The diagnostic accuracy indices of both the tests are given in table 5. Negative predictive value (NPV) for both ST IgM ICT and the ST IgM ELISA is >99% implying that a negative result is definitely not a case of scrub typhus

**Table 5:** Diagnostic accuracy of ST IgM ELISA and ST IgM ICT (95% CI)

| Diagnostic accuracy | ST IgM ICT(95% CI) | ST IgM ELISA |
| --- | --- | --- |
| Sensitivity | 98.7(93,100) | 97.4(90.9,99.7) |
| Specificity | 96.3(93.3,98.2) | 95.9(92.8,97.9) |
| LR+ | 26.6(14.4,48.8) | 23.8(13.3,42.5) |
| LR- | 0.0135(0.0019,0.0946) | 0.0271(0.0069,0.106) |
| PPV | 88.4(79.7,94.3) | 87.2(78.3,93.4) |
| NPV | 99.6(97.9,100) | 99.2(97.2,99.9) |

IgM ELISA test OD values were studied separately. ROC curve was constructed with OD values to discriminate the diseased from the non-diseased and the same is represented in Fig 1

**Figure 1:**
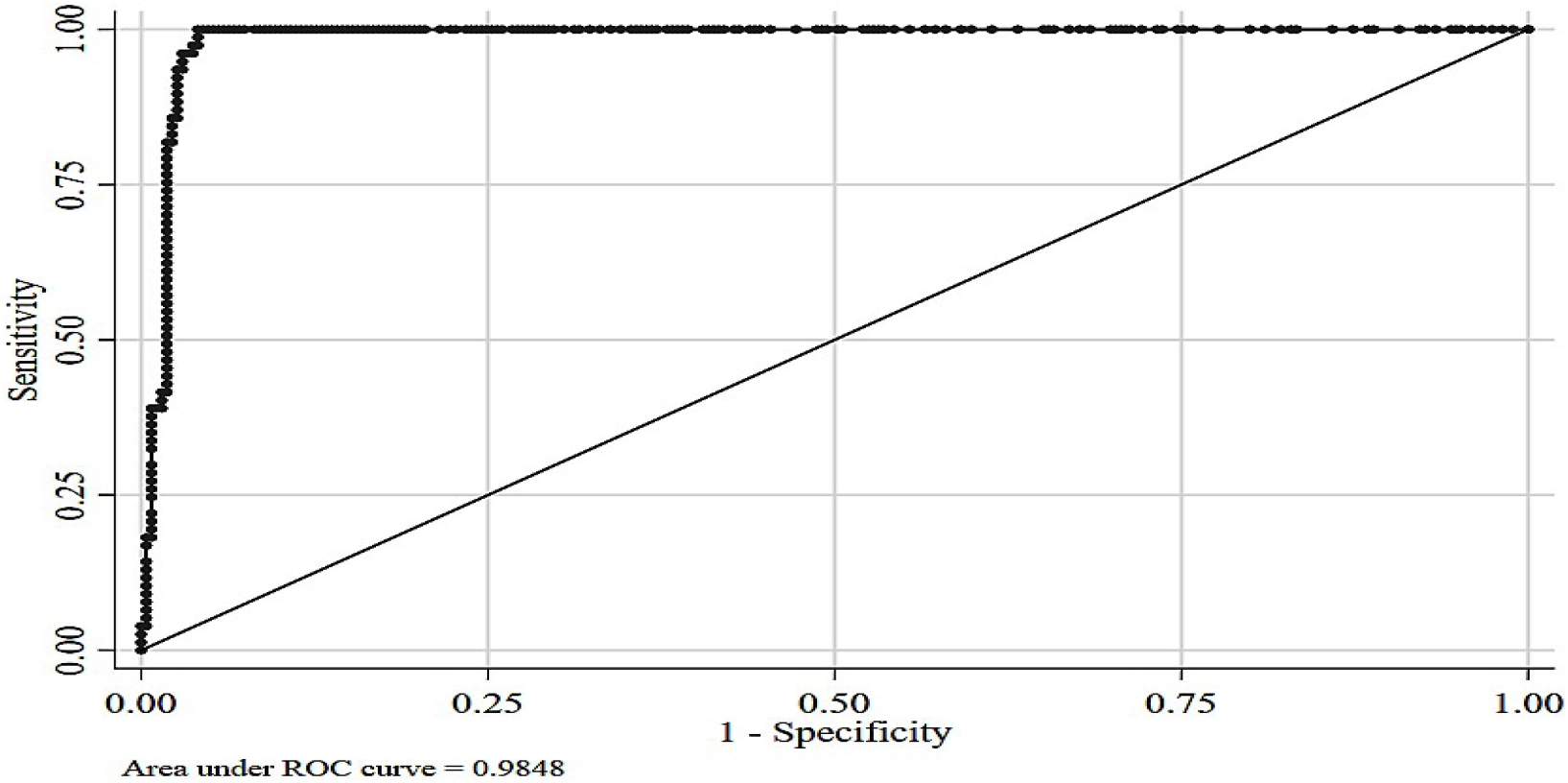
ROC curve for ST IgM ELISA.

Receiver operating characteristic curve (ROC) for the ST IgM ELISA performed on a serum dilution of 1:100 at a diagnostic cutoff OD ≥ 1.0 is shown in fig 1. Area under the curve using this serum dilution and diagnostic cutoff falls in excellent category (0.9 – 1.0) as it is very close to 1.0.

The performance indices of ELISA at different OD cut-off values are given in Table 6. It shows that the best sensitivity and specificity and correct classification as a case of scrub typhus occurred at an OD of 1.0 ^(16)^(serum dilution is 1:100)

**Table 6:**
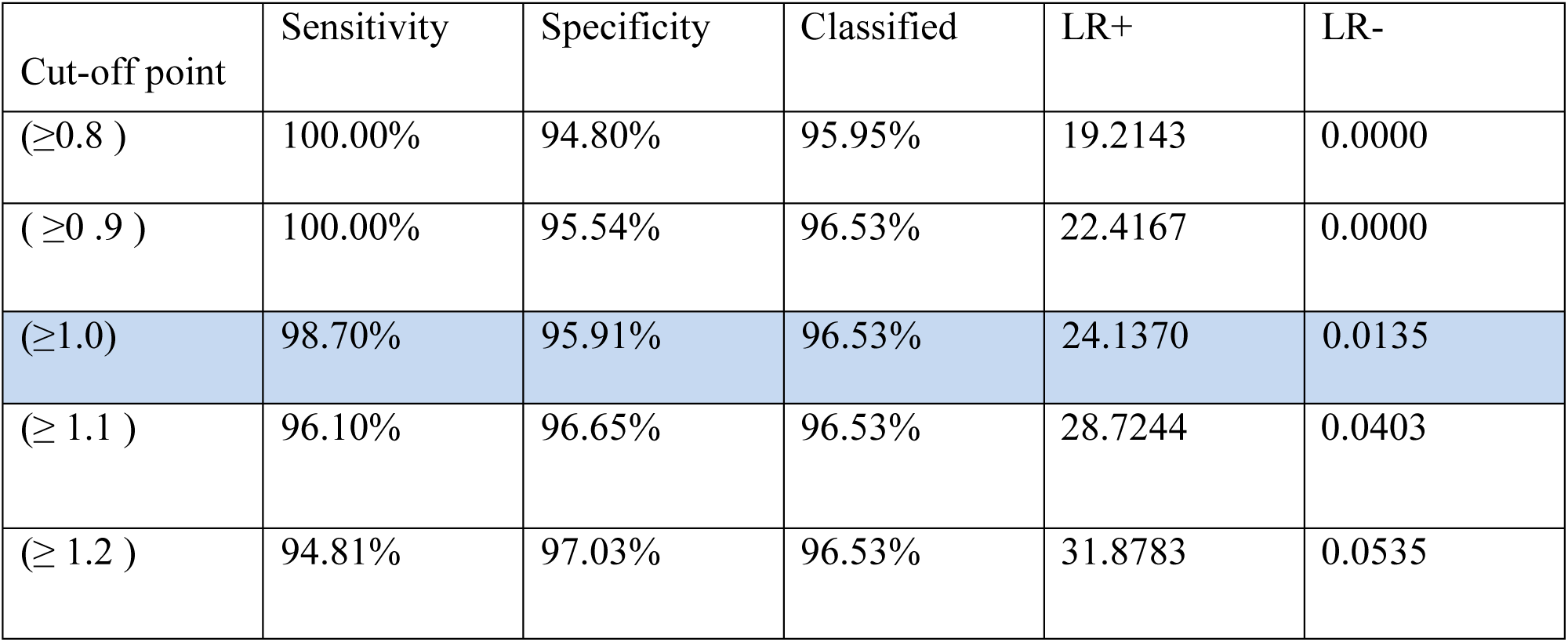
Performance indices of ST IgM ELISA at different cut off values (Based on ROC analysis)

| Cut-off point | Sensitivity | Specificity | Classified | LR+ | LR- |
| --- | --- | --- | --- | --- | --- |
| ($\geq 0.8$) | 100.00% | 94.80% | 95.95% | 19.2143 | 0.0000 |
| ($\geq 0.9$) | 100.00% | 95.54% | 96.53% | 22.4167 | 0.0000 |
| ($\geq 1.0$) | 98.70% | 95.91% | 96.53% | 24.1370 | 0.0135 |
| ($\geq 1.1$) | 96.10% | 96.65% | 96.53% | 28.7244 | 0.0403 |
| ($\geq 1.2$) | 94.81% | 97.03% | 96.53% | 31.8783 | 0.0535 |

## Discussion

The need for technical expertise and costs incurred have turned our attention to exploring the role of RDTs in diagnosing ST in resource-limited settings.^1^ In this study, well-characterized sera from normals and patients with scrub typhus, malaria, dengue, enteric fever and gram negative septicaemia which are common causes of AFI have been included. Cross reactions have been seen in dengue positive sera and sera belonging to patients who were smear positive for malarial parasites. While two of the six cases with positive dengue serology had an eschar and demonstrated defervescence of fever within 48 hours of initiation of doxycycline therapy, two others did not have an eschar but responded to doxycycline. Therefore these four cases were considered definite scrub typhus cases as per a composite case definition (data from IJMM, accepted for publication in press) derived at the study center. The remaining two cases were not considered as true scrub typhus cases as both had no eschar and showed no response to ST-specific therapy. The number of false positives picked up in smear confirmed cases of malaria were less using IgM ICT than with IgM ELISA. These malaria positive serum samples were tested for assessing the analytical specificity of scrub typhus IgM ELISA as previous studies have shown cross reactions.^17,18,19^ The high negative predictive value (>99%) of both the scrub typhus IgM ICT and IgM ELISA including the excellent concordance (Kappa value 0.97) suggests that IgM ICT can be used interchangeably with IgM ELISA to rule out scrub typhus. Furthermore, the ICT will be an excellent rapid diagnostic test in emergency departments, and primary health care centers

Determining the grey zone in ELISA tests and repeat testing of the samples falling in this zone will enhance the discriminatory performance of the test as has been demonstrated by Solanki et al.^20^ Based on the performance indices, OD value of 1.0 ±0.2 on ELISA need to be retested or verified with a rapid test. If on repeat testing the grey zone sample showed one or both OD values above the cut-off value it was marked as ‘reactive’in classification.

A systematic review and meta-analysis on point of care testing in scrub typhus by Saraswati *et al* showed that the sensitivity and specificity of various ICTs ranged from 23.3% to 100% and 73% to 100% respectively.^21^ Studies from India on RDTs as serological tests for diagnosis of ST are very few. The diagnostic accuracy indices obtained in our study are similar to those by Anita Raj *et al* who also evaluated InBios Scrub TyphusDetect IgM Rapid Test with InBios Scrub Typhus Detect IgM ELISA.^22^ But our study had well characterized sera which could elaborate on cross reactions giving rise to false positive test results. Pote *et al* used immunochromatographic test (RDT) which showed low sensitivity but high specificity.^11^ Again, well characterized sera and other indices like PPV and NPV were not mentioned. Though ImmuneMed Scrub Typhus Rapid, another RDT showed good sensitivity, specificity, PPV and NPV^22,23,24^ the kit is not yet available in India. Kingston *et al* used well characterized samples to determine optimal RDT performance with antibody titre in IFA as comparator.^25^ The results showed good sensitivity and specificity at high IFA titres and is therefore expected to perform well in endemic settings. It holds good in our setup with the high prevalence of scrub typhus showing a seasonal trend.

There is a definite need to develop representative and well-characterized scrub typhus endemic and non-endemic cohorts from other regions to evaluate the diagnostic capacity of new and existing assays.

The limitations of this study include not testing the sera for leptospirosis but this is of not much consequence as this disease is very uncommon amongst patients presenting with AFI to our centre(since it is hardly ever seen in our study population)^26^. Lack of using samples confirmed by molecular assays is another drawback as this would have helped in resolving false positive results.

The strength of this study lies in inclusion of prospectively collected, well-characterized sera from patients with acute undifferentiated febrile illness and determining the accuracy indices appropriately while explaining the cross reactions.

## Conclusion

The availability of infrastructure and capacity for accurate diagnosis of scrub typhus is not uniform in distribution throughout the health care settings in the country. The costs incurred and expertise required limit the access to standard serological tests. The need for rapid diagnostic tests with their simplicity and speed in resource limited settings where most of the cases occur, cannot be overemphasized. The findings from this study suggest that IgM ICT can substitute IgM ELISA in such settings. Further prospective studies from different centres in India are needed to confirm the same.

